# Localizing the chaperone activity of erythroid spectrin

**DOI:** 10.1101/534982

**Authors:** Dipayan Bose, Abhijit Chakrabarti

## Abstract

Spectrin, the major protein of the RBC membrane skeleton has canonically been thought to only serve a structural function. We have described a novel chaperone-like property of spectrin and have shown that it is able to prevent the aggregation of other proteins such as alcohol dehydrogenase, insulin and free globin chains. We have tried to localize the molecular origin of chaperone-like activity in multi-domain spectrin by using recombinant spectrin fragments and investigating individual domains. We have characterized the recombinant domains using intrinsic tryptophan fluorescence and CD spectroscopy to show their identity to native spectrin. Hydrophobic ligands Prodan (6-propionyl-2[dimethylamino]-naphthalene) and ANS (1-anilinonaphthalene-8-sulfonic acid) binding has been used to probe the hydrophobicity of the recombinant domains and it is seen that all domains have surface exposed hydrophobic patches; and in accordance with our previous hypothesis only the reconstituted self-association domain binds Prodan. Recombinant domains display comparable chaperone potential in preventing protein aggregation; and substrate selectivity of α-over β-globin is seen. Enzyme refolding studies show alternate pathways of chaperone action. Our current study points to the presence of hydrophobic patches on the surface of these domains as the source of the chaperone activity of spectrin, as notably seen in the self-association domain. There is no one domain largely responsible for the chaperone activity of spectrin; rather all domains appear to contribute equally, such that the chaperone activity of spectrin seems to be a linear sum of the individual activities of the domains.

## Introduction

Spectrin is a protein that is found in the membrane skeleton of all metazoan cells (1,2). It is the major scaffolding component of mature erythrocytes in humans (3). It is large (∼500 kDa), heterodimeric protein composed of α (∼240 kDa) and β (∼220 kDa) polypeptides (4). Each polypeptide subunit is composed mostly of tandemly repeating spectrin repeat domains which are 106 amino acid long stretches that fold into three anti-parallel α-helices (5). The spectrin repeat domains have been implicated as the source of the remarkable elasticity, flexibility and structural strength of spectrin (6-8). They have also been shown to be the location of the protein binding and ligand binding property of spectrin (9,10). These domains have a high degree of homology (11), especially the presence of conserved tryptophans (12), within themselves; however they also have sufficiently different sequences to show distinct properties (13). There are 20 such repeat domains in α-spectrin and 16 domains in β-spectrin and the rest of the polypeptide sequences are made up of non-homologous motifs (4,14).

Previous literature on spectrin mostly points out the role of spectrin as a structural and mechanical component of the membrane skeleton (3,15,16). Newer studies point out the non-canonical roles of spectrin, for example as a platform for signal transducing complexes and as a protein-protein interaction module (13,17,18). The first report of the chaperone like property of erythroid spectrin was made by our group where we had also hypothesised that this property resides in the self-association domain of the hetero-dimer (19-22).

Keeping in mind the modular, repeating nature of the spectrin repeat domains of which the majority of erythroid spectrin is constituted it becomes important to determine if the chaperone property of spectrin is localized in one of these repeat domains or is a shared general property of these repeat motifs or is a function of the non-homologous motifs.

Spectrin presents an interesting case where like many cytoskeletal proteins, it was believed that the spectrin repeat domains serve only to flexibly link and spatially separate actin-binding function (23,24); however it has now been shown that the repeat domains too have their own functionality. Moreover the interactions of these repeat domains cannot be generalized even though they share sequence and structural identity (10,13,25). Thus, keeping our question in mind, it becomes difficult to predict the functionality of a given repeat. Moreover spectrin has known functionality in some of its repeat domains such as actin binding and ankyrin binding (26,27), and there is homology with known chaperone calmodulin in its EF domain (28). This further complicates the question of localizing spectrin’s chaperone activity. In our current work we have tried to test our old hypothesis and determine if the chaperone like potential of spectrin is localized in one of its many domains or is a property displayed by the entire polypeptide.

It is known that all chaperones have some features in common which aid in substrate recognition; most often these are charged and/or hydrophobic patches which are solvent exposed (29). Keeping in mind the known presence of hydrophobic patches on spectrin as well as our hypothesis linking hydrophobic patches to chaperone activity (20,30), the most likely loci on spectrin to harbour chaperone activity should also have hydrophobic patches. In our present study we have selected nine domains of spectrin, five from α and four from β spectrin to investigate their chaperone activity as versus the native protein. We have selected domains based on their likelihood of harbouring hydrophobic patches/chaperone like activity and the domains we have chosen to investigate are the α-tetramerization domain, α-dimerization domain, EF domain, SH3 domain, β-tetramerization domain, β-dimerization domain, actin binding domain, ankyrin binding domain, and a random non-specific spectrin repeat domain from α-spectrin.

The self association domain or Tetramerization domain of spectrin is the dimerized form of the N-terminal 1-158 amino acids of α (α-tetramerization domain) and C-terminal 1895-2137 amino acids of β-spectrin (β-tetramerization domain) (31). We have previously hypothesized that the chaperone function and Prodan binding site of spectrin is localized here (20). Available literature indicates that this self association domain is where the hetero-dimers of spectrin link up to form tetramers (32) and mutations in this region lead to haemolytic diseases (33). Moreover available data shows this region to have hydrophobic residues present on the peptide surface (34,35). This presence of surface exposed hydrophobic patches has led us to include these two domains.

Similarly, we have selected the α-dimerization domain (2002-2233 amino acids of α-spectrin) and β-dimerization (271-500 amino acids of β-spectrin) domain of spectrin, which are the regions in the polypeptides that form the nucleation site of high affinity hetero-dimer formation and also have surface hydrophobic patches (36-39).

The C-terminal of α-spectrin is calmodulin like and is called the EF domain (2257-2429 amino acids) and has calcium dependent and independent EF hands; since calmodulin itself is a chaperone this domain too is a candidate for possessing chaperone like function (28,40).

SH3 domain (973-1055 amino acids of α-spectrin) is a protein interaction module present in α-spectrin and due to its broad range of protein interaction ability is also considered (41,42).

The ankyrin binding domain (1657-1876 amino acids of β-spectrin) (26,43) and actin binding domain (1-316 amino acids of β-spectrin) (27,44,45) of the of β-spectrin polypeptide are also protein binding domains and are thus included.

As versus these domains which have some defined function and probable chaperone potential a random spectrin repeat domain of α-spectrin between amino acids 1470-1576 was chosen to act as a control (4).

We have expressed and purified these chosen domains from cloned sequences and characterized them using intrinsic tryptophan fluorescence, urea denaturation, CD-spectroscopy, time-resolved spectroscopy and ligand binding. We have estimated their chaperone potential by monitoring the extent of protection of protein aggregation thermally and non-thermally and by enzyme refolding assays.

## Results

### Expression and characterization of spectrin domains

The spectrin domains were expressed, extracted and purified in urea denatured condition, and then refolded to give native domains. The domains of spectrin that were selected for the present study are pictorially depicted in **Figure 1,** panel 1 for easy visualization of their relative locations in spectrin. The spectrin domains were expressed and purified as described and was run on 15% SDS-PAGE to check purity; **Figure 1** panel 2, subpanels ‘a’ and ‘b’ show gels for crude and purified extracts respectively. Supplementary **Table ST1** contains the polypeptide sequence for each domain and their calculated molecular weight. **Figure 1** panel 4 shows the fluorescence emission spectra of the individual native fragments and reconstituted self-association domain. Supplementary **Figure S1** shows the same for urea denatured condition. It is seen that for most of the domains the emission maxima is close to that of intact dimeric spectrin and like the parent protein there is red shifting of emission upon urea unfolding. The parameters of fluorescence lifetime, anisotropy and polarization are tabulated in **Table T1**. It is seen that for almost all the fragments the parameters are close to spectrin. The self-association domain was reconstituted from α and β-tetramerization domains and was run on a Sephadex G-100 column for purification. It was seen that over a 12 hour period the majority of constituent fragments had dimerized to form the self-association domain. The elution profile for the same is given in **Figure 1** panel 3. CD spectroscopy reveals that the native domains all have an α-helical fold analogous to spectrin, except the SH3 domain which shows a more β-sheet like nature with the presence of a unique secondary minima which matches with previous literature (46). The CD spectra of spectrin superposed with that of the self-association domain is shown in **Figure 1** panel 5 and that of the rest of the fragments are shown in supplementary **Figure S2**. The MRE values at 222 nm are tabulated in **Table T1.** Upon urea denaturation it was seen that the domains lost most of their secondary structure unlike spectrin which still retained some of its secondary structure. Representative CD spectra of the urea denatured domains are shown in supplementary **Figure S3.**

**Figure 1:**
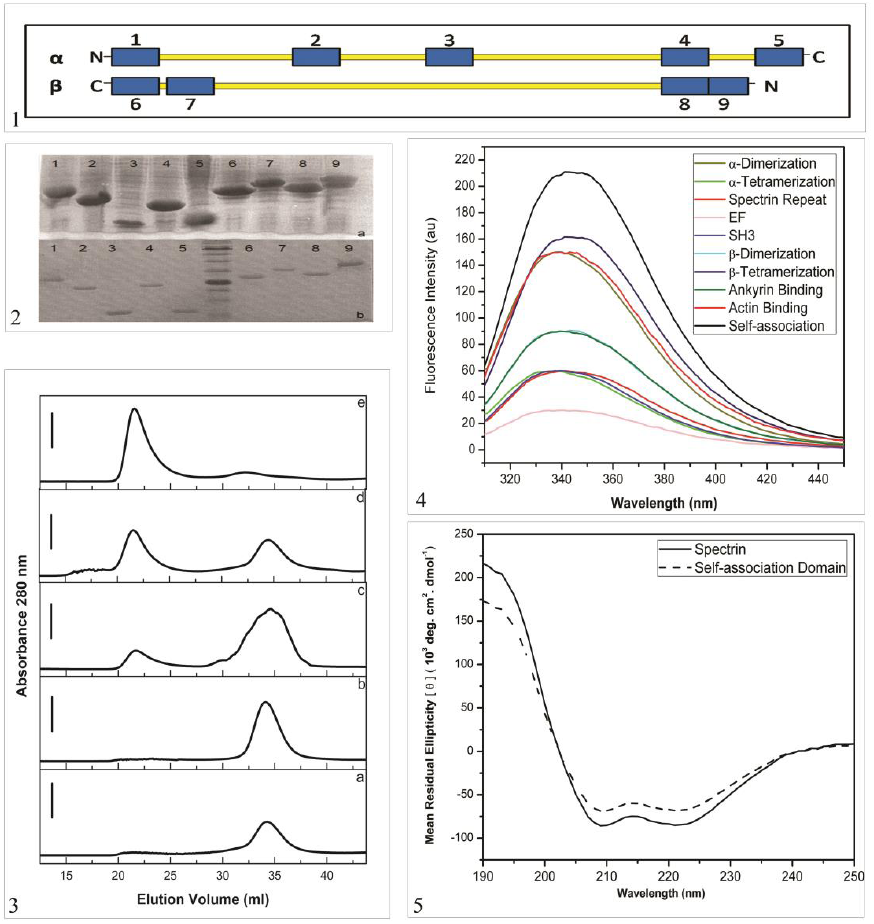
Panel 1 shows a pictorial representation of the spectrin domains chosen for this study and their location in dimeric spectrin. The anti-parallel spectrin dimer is shown in yellow and the domains are highlighted in blue. The domains are numbered 1 through 9 and in numerical sequence they are: α-tetramerization domain, SH3 domain, spectrin repeat domain, α-dimerization domain, EF domain, actin binding domain, β-dimerization domain, ankyrin binding domain and β-tetramerization domain respectively. Panel 2 subpanel-a shows the SDS-PAGE for the crude extracts of the domains and subpanel-b shows the same for Ni-NTA resin purified samples. The lanes are numbered from 1 through 9 and in numerical sequence they are: α-dimerization domain, α-tetramerization domain, spectrin repeat domain, EF domain, SH3 domain, β-dimerization domain, β-tetramerization, ankyrin binding domain and actin binding domain respectively. The ladder used was Prism Ultra pre-stained protein ladder and the two most prominent bands show 24 kDa and 70 kDa. Panel 3 shows the reconstitution of the self-association domain from the α- and β-tetramerization domains as followed by size exclusion chromatography on Sephadex G-100 column. Subpanels a and b show the elution profile for purified α-(∼19 kDa) and β-tetramerization (∼28 kDa) domains respectively. Subpanel-c shows the same for when the two domains are mixed and incubated for two hours, it is seen that a small self-association domain (∼47kDa) peak starts to form. Subpanel-d shows increased self-association domain production at 4 hours and subpanel-e shows that the majority of the constituent domains have dimerized to form self-association domain at 8 hours. Panel 4 shows the emission spectra of all the native spectrin domains, spectrin and self-association domain while panel 5 shows the representative CD spectra of native spectrin and reconstituted self-association domain.

### Binding of fluorescent ligands

The emission maxima of Prodan in aqueous solution was found to be blue shifted from 520 nm to 430 nm upon spectrin and self-association domain binding with a concurrent large increase in emission intensity. It was found that other than the reconstituted self-association domain other spectrin domains did not appreciably bind with Prodan to a level that could be experimentally followed within the concentration range of the soluble spectrin fragments. Prodan binding was analysed using both the model independent and Scatchard methods. The dissociation constants and stoichiometries derived from these two methods are tabulated in **Table 2**. It was also noted that upon binding the fluorescence lifetime, polarization and anisotropy of Prodan increased appreciably; data is given in **Table 2**. The representative fluorescence spectra and binding isotherms of Prodan binding to self-association domain are shown in **Figure 2**.

**Table 1:** The emission maxima for 295 nm excitation; anisotropy, polarization and mean lifetime values for 295 nm excitation, 340 nm emission, of the tryptophans of the native spectrin domains and spectrin are tabulated. Values in parenthesis indicate the same for 8 M urea denatured condition. The α-helical content is represented as the mean residual ellipticity at 222 nm. Error values are calculated as the standard deviation of four independent experiments.

| Domain/<br>protein | $\lambda_{\max}$ (ex.<br>295nm) | Anisotropy (r) | Polarization (p) | Mean<br>Lifetime<br>(n.s.) | Mean<br>Residual<br>Ellipticity<br>[$\theta$] ($10^3$<br>deg. cm<sup>2</sup>.<br>dmol<sup>-1</sup>) |
| --- | --- | --- | --- | --- | --- |
| α-<br>Tetramerization | 336 ± 1.2<br>(347 ± 1.3) | 0.083 ± 0.002<br>(0.054 ± 0.001) | 0.121 ± 0.003<br>(0.067 ± 0.004) | 3.87 ± 0.2<br>(3 ± 0.1) | -110.6 ±<br>2.2 |
| SH3 | 339 ± 1.1<br>(347 ± 1.1) | 0.072 ± 0.001<br>(0.044 ± 0.002) | 0.121 ± 0.002<br>(0.06 ± 0.003) | 3.5 ± 0.3<br>(2.9 ± 0.1) | -17.5 ± 5.3 |
| Spectrin<br>repeat | 342 ± 1.4<br>(347 ± 1.5) | 0.073 ± 0.001<br>(0.034 ± 0.002) | 0.124 ± 0.003<br>(0.045 ± 0.003) | 3.44 ± 0.2<br>(2.7 ± 0.3) | -134.1 ±<br>3.1 |
| α-<br>Dimerization | 337 ± 1.1<br>(346 ± 1.1) | 0.087 ± 0.002<br>(0.041 ± 0.001) | 0.101 ± 0.003<br>(0.05 ± 0.002) | 3.7 ± 0.1<br>(3.1 ± 0.1) | -118.6 ±<br>1.1 |
| EF | 340 ± 1.2<br>(348 ± 1.3) | 0.063 ± 0.001<br>(0.042 ± 0.001) | 0.09 ± 0.003<br>(0.056 ± 0.003) | 3.6 ± 0.2<br>(2.9 ± 0.2) | -59.3 ± 4.3 |
| Actin Binding | 337 ± 1.1<br>(343 ± 1.1) | 0.083 ± 0.002<br>(0.045 ± 0.001) | 0.117 ± 0.002<br>(0.063 ± 0.002) | 4.0 ± 0.3<br>(3.28 ± 0.3) | -105 ± 2.2 |
| β-<br>Dimerization | 337 ± 1.2<br>(344 ± 1.3) | 0.082 ± 0.002<br>(0.05 ± 0.001) | 0.12 ± 0.003<br>(0.065 ± 0.001) | 4.1 ± 0.1<br>(3.5 ± 0.1) | -97.5 ± 4.6 |
| Ankyrin<br>Binding | 336 ± 1.1<br>(348 ± 1.2) | 0.092 ± 0.001<br>(0.041 ± 0.001) | 0.148 ± 0.002<br>(0.073 ± 0.002) | 3.8 ± 0.2<br>(3.49 ± 0.3) | -93.5 ± 3.2 |
| β-<br>Tetramerization | 338 ± 1.1<br>(346 ± 1.1) | 0.086 ± 0.002<br>(0.044 ± 0.001) | 0.113 ± 0.002<br>(0.051 ± 0.003) | 3.9 ± 0.2<br>(3.3 ± 0.3) | -49.5 ± 4.4 |
| Spectrin | 338 ± 1.1<br>(345 ± 1.1) | 0.12 ± 0.001<br>(0.04 ± 0.001) | 0.18 ± 0.002 (0.04<br>± 0.002) | 3.8 ± 0.1<br>(2.9 ± 0.1) | -84.8 ± 3.3 |
| Self-association<br>domain | 337 ± 1.2<br>(346 ± 1.3) | 0.1 ± 0.002<br>(0.06 ± 0.001) | 0.147 ± 0.003<br>(0.076 ± 0.002) | 3.9 ± 0.2<br>(2.8 ± 0.3) | -67.5 ± 4.1 |

**Table 2:** The K_d_ values and stoichiometry of spectrin and self-association domain binding to Prodan are shown as derived from the model independent method. K_0_ values and stoichiometry are shown in parenthesis as derived from the Scatchard plot. The anisotropy, polarization and mean lifetimes of Prodan bound to spectrin and self-association domain are tabulated for 360 nm excitation and 430 nm emission. Values in parenthesis represent the same for Prodan in aqueous buffer. Error values are calculated as the standard deviation of four independent experiments.

| Domain/<br>Protein | $K_d/K_0$ value<br>($\mu\text{M}$) | Stoichiometry<br>(n) | Anisotropy<br>(r) | Polarization<br>(p) | Fluorescence<br>Lifetime (ns) |
| --- | --- | --- | --- | --- | --- |
| Spectrin | $0.44 \pm 0.1$<br>($0.38 \pm 0.13$) | $1.1 \pm 0.1$<br>($1 \pm 0.1$) | $0.18 \pm 0.001$<br>($0.02 \pm 0.001$) | $0.28 \pm 0.001$<br>($0.03 \pm 0.001$) | $5.4 \pm 0.3$<br>($1.5 \pm 0.2$) |
| Self-association<br>Domain | $14.3 \pm 0.3$<br>($12.5 \pm 0.4$) | $1 \pm 0.2$<br>($1.1 \pm 0.1$) | $0.16 \pm 0.002$ | $0.27 \pm 0.002$ | $5.2 \pm 0.4$ |

**Figure 2:**
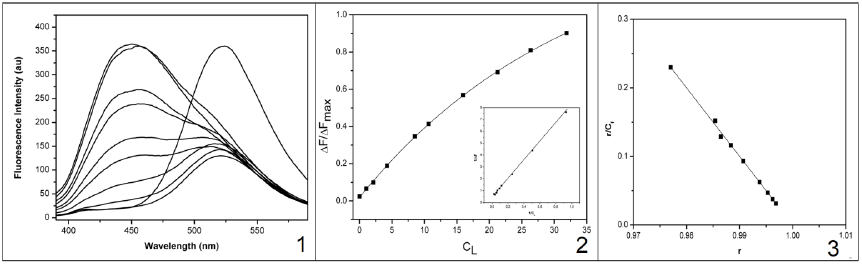
Panel 1 shows the fluorescence spectra of Prodan titrated with increasing concentration of the self association domain. The excitation was at 360 nm, unbound emission was at 520 nm and bound emission at 430 nm. Sequential decrease in un-bound and increase of bound emission peak was seen on titrating with spectrin. Panel 2 shows the binding isotherms calculated by the model independent method and panel 3 shows the same for that calculated by the Scatchard method.

Similarly in case of ANS it was seen that the fluorescence emission maxima was found to be blue shifted from 470 nm to 520 nm upon protein binding and the emission intensity was found to increase many fold over that of free ANS in aqueous buffer which has negligible fluorescence. The binding parameters of ANS to the spectrin fragments were evaluated by two methods as before and the stoichiometries and binding constants are tabulated in **Table 3**. The increase in fluorescence upon self-association domain binding, and the analysis are graphically represented in **Figure 3**. The same for the rest of the domains are shown in supplementary **Figures S4** and **S5**. The increase in polarization and anisotropy of ANS fluorescence upon protein binding are tabulated in **Table 3**.

**Table 3:**
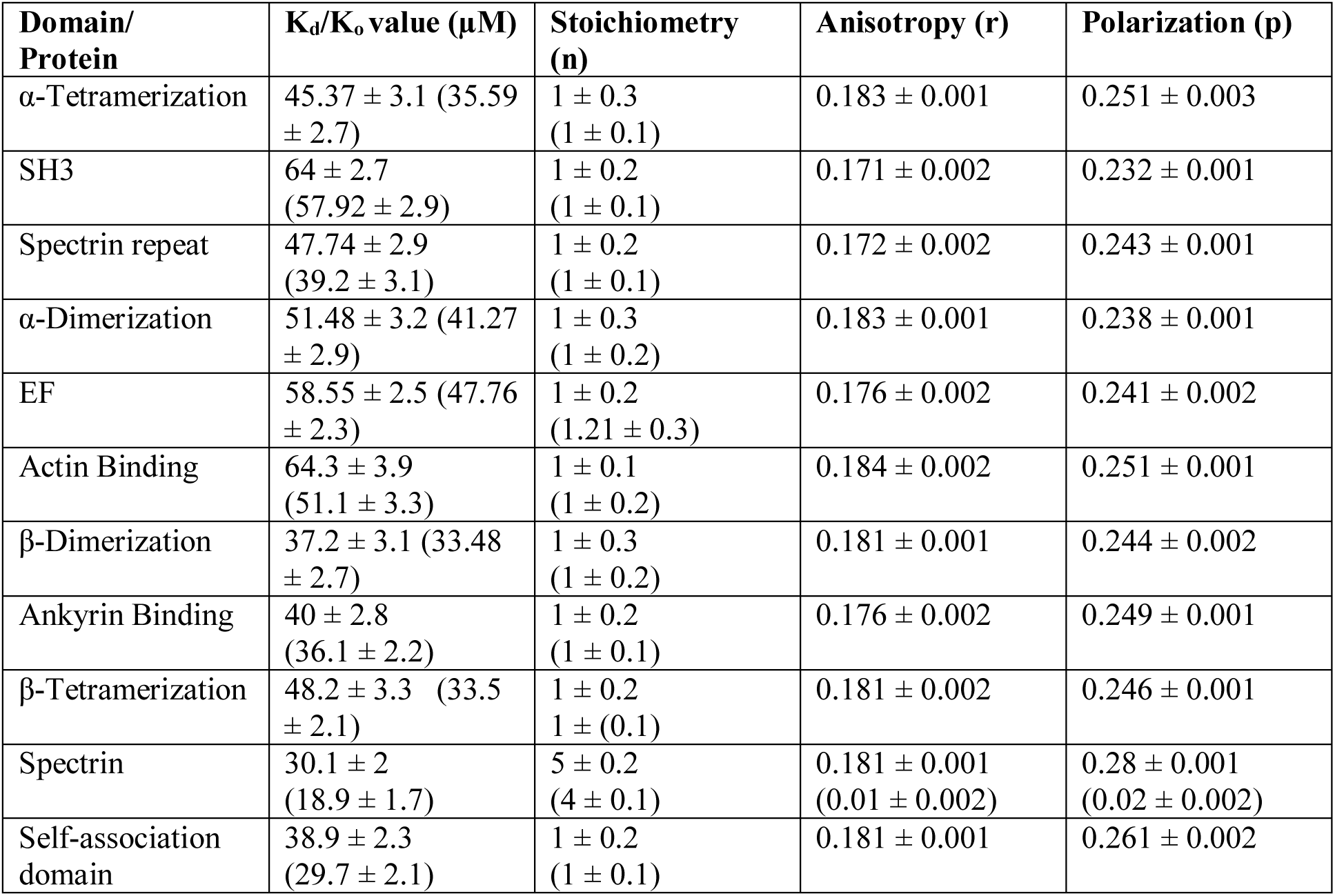
The K_d_ values and stoichiometry of spectrin, self-association domain and spectrin domain binding to ANS are shown as derived from the model independent method. K_0_values and stoichiometry are shown in parenthesis as derived from the Scatchard plot. The anisotropy, polarization and mean lifetimes of ANS bound to spectrin, self-association domain and spectrin domains are tabulated for 372 nm excitation and 470 nm emission. Values in parenthesis represent the same for ANS in aqueous buffer. Error values are calculated as the standard deviation of four independent experiments.

**Figure 3:**
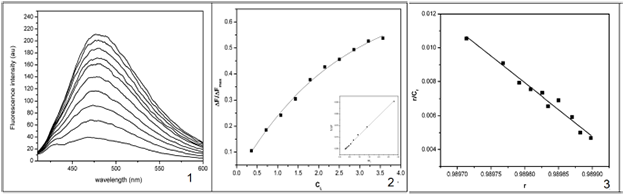
Panel 1 shows the fluorescence spectra of self-association domain titrated with increasing concentration of the ANS. The excitation was at 372 nm, and bound emission at 470 nm. Sequential increase of bound emission peak was seen on titration, with unbound ANS having negligible fluorescence. Panel 2 shows the binding isotherms calculated by the model independent method and panel 3 shows the same for that calculated by the Scatchard method.

### Assay of chaperone activity

In case of protein aggregation the maximum O.D. measured of the test proteins aggregating alone was taken as 100% and the reduction thereof by spectrin and its cloned fragments was expressed as the percentage of protection from aggregation. It was seen that spectrin and spectrin fragments as well as the reconstituted self-association domain could all inhibit protein aggregation both in an aggregating protein to chaperone ratio of 1:0.5 and 1:1. It was seen that this inhibition of aggregation was dose dependent where more protection occurred for the 1:1 ratio than for the 1:0.5 ratio. Moreover it was seen that in case of insulin and globin chain aggregation the fragments behaved in a general way showing a similar extent of protection for all cases with the extent of protection being only slightly less than that of intact spectrin. In case of the thermal aggregation of ADH it was seen that the fragments showed a decreased overall protection than in the other two cases. For the aggregation of insulin native spectrin showed a protection of ∼45 % in 1:0.5 weight ratio and about ∼60 % for the 1:1 weight ratios. The fragments all showed a protection of ∼ 35-40% and 50-55% for these two ratios respectively. In case of ADH aggregation, native spectrin showed ∼ 40 % protection in the 1:0.5 and ∼ 50% protection in the 1:1 weight ratio respectively with the fragments showing ∼ 30 % and ∼ 40 % protection in these two weight ratios respectively. For the globin chains it was seen that both native spectrin and the fragments were able to better protect α-globin from aggregation rather than β-globin. Representative aggregation curves of insulin aggregating n presence and absence of spectrin and its subunits is shown in **Figure 4. Figure 4** also shows the extent of protection from aggregation of the test proteins insulin, ADH, α and β-globin by spectrin and its fragments.

**Figure 4:**
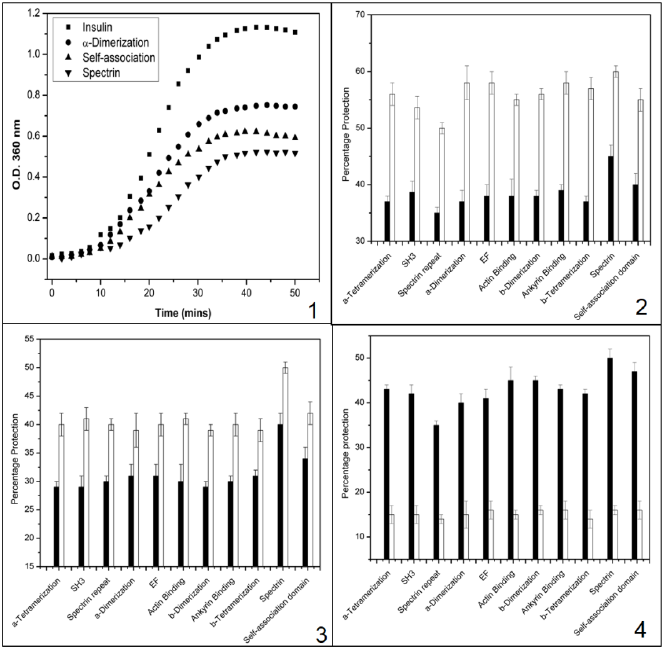
Panel 1 shows representative aggregation curves of insulin aggregating alone or in presence of spectrin, reconstituted self-association domain or α-dimerization domain in a insulin: chaperone ratio of 1:0.5. Aggregation was followed by monitoring the apparent increase in absorbance at 360 nm. Panel 2 shows the extent of protection as percentage from aggregation of insulin by spectrin, spectrin domains and self-association domain where the aggregation of insulin alone is considered as 100%. Filled in bars represent protection for 1:0.5 weight ratio and empty bars represent the same for 1:1 weight ratio. Panel 3 shows the extent of protection as percentage from aggregation of ADH by spectrin, spectrin domains and self-association domain where the aggregation of ADH alone is considered as 100%. Aggregation was followed by measuring the 90° scattering at 450 nm. Filled in bars represent protection for 1:0.5 weight ratio and empty bars represent the same for 1:1 weight ratio. Panel 4 shows the percentage protection against aggregation of α- and β-globin by spectrin, self-association domain and spectrin domains. The filled in bars represent protection in case of α-globin and the empty bars represent that of β-globin. Aggregation was followed by measuring apparent increase in absorbance at 700 nm and the extent of aggregation of either of the two globins aggregating alone was taken to be 100%. Error bars are calculated as the standard deviation of four independent experiments.

It was seen that native spectrin and its recombinant fragments had an effect on the reactivation yield of the enzymes α-glucosidase and alkaline phosphatase. In case of alkaline phosphatase it was seen that presence of spectrin or its fragments caused an increase in the reactivation yield whereas in case of α-glucosidase there was a decrease. In case of alkaline phosphatase it was seen that native spectrin could increase the yield to ∼35% from a self refolding yield of ∼ 20%. The fragments could increase the yield comparably to spectrin at around 30%. In case of α-glucosidase self refolding yields of ∼25% was found to decrease in presence of spectrin to ∼ 10% with the around same yield being seen in presence of the fragments. The reactivation yields are graphically represented in **Figure 5.**

**Figure 5:**
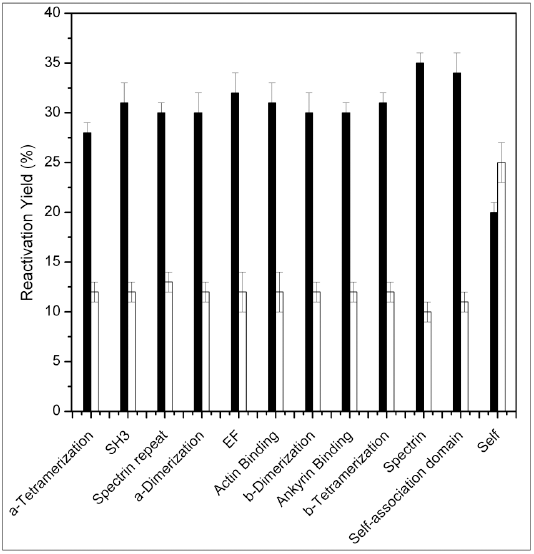
The percentage reactivation yields of alkaline phosphatase and α-glucosidase are shown. The filled in bars represent alkaline phosphatase while the empty bars represent α-glucosidase. The activity of equivalent amount of native enzyme was taken as 100%. Presence of spectrin, self-association domain and spectrin fragments was found to reduce reactivation yields for α-glucosidase, with the opposite being seen for alkaline phosphatase. Error bars are calculated as the standard deviation of four independent experiments.

## Discussions

Intrinsic tryptophan fluorescence is a very useful tool to probe the conformation and stability of spectrin. The tryptophans are conserved in their positions in the spectrin repeat domains and contribute to the overall stability and structure of the protein as evidenced by the fact that mutating them causes a decrease in stability of the spectrin repeat domains (12,47,48). Of our chosen spectrin domains all but the SH3, EF, Actin binding and the β-tetramerization domain are made up of the spectrin repeat motifs, and of the domains that do not share a consensus sequence with the spectrin repeat motif, only the SH3 domain has a non-alpha helical fold (49). So tryptophan fluorescence gives us a useful tool to probe the structure of the cloned spectrin domains. Experimental data shows that for all the domains a moderately high amount of constraint is present on the tryptophans, which is reflected by their anisotropy and polarization values which approach those of native spectrin (50,51). Notably, the SH3 domain and EF domain are exceptions to that general trend which can be explained by the fact that SH3 domain has a globular fold which does not put as much torsional constraint on the tryptophans and an in case of the EF domain the single tryptophan being probed was inserted artificially to aid in estimating protein folding. The anisotropy data is corroborated by emission maxima measurements which show the same trend and it is seen that the tryptophan emissions corroborate well with situations where the tryptophans are buried in a hydrophobic protein core (52). Moreover upon urea denaturation the anisotropy decreases and emission maxima are red shifted for the refolded domains as it is for native spectrin (53). Taken together the anisotropy, polarization and emission maxima data strongly indicate that the cloned, expressed and denatured domains were successfully refolded and the refolded proteins were structurally similar to native spectrin. This is supported by measurements of the mean tryptophan lifetimes which show that the refolded domains all have lifetimes comparable to that of native spectrin which decreases upon urea unfolding analogous to spectrin. CD spectroscopy confirmed the folding of the proteins; it was seen that the refolded domains shared the same α-helical fold of spectrin except SH3 domain which showed a CD spectra that matched with previous reports of the native domain (46). Urea denaturation was found to abolish the clean α-helical nature of the CD spectra of these domains further indicating that the domains were successfully renatured.

Interestingly it is seen that CD spectra reveals subtle differences in the folding and conformation of these spectrin domains which are consistent with previous literature on these domains. CD spectroscopy reveals that the lowest MRE values at 222 nm are those of the EF domain, SH3 domain and β-tetramerization domain, which can be explained by the fact that β-dimerization domain has a non-structured tail (54) which decreases its α-helical nature; similarly SH3 domain has a globular fold (46) and EF domain while α-helical is made up sequences non-homologous to the spectrin repeat domain (28).

Hydrophobic probe Prodan was found to bind only the reconstituted self-association domain of spectrin with binding affinity comparable to native spectrin. The rest of the fragments either do not bind Prodan or bind weakly with appreciably lower binding affinity. This validates our earlier hypothesis that the Prodan binding site in spectrin is located in the self-association domain; moreover the stoichiometry is also found to be same as that seen in our previous studies (20,55). It is important to note that our previous study has shown that Prodan binds to both dimeric and tetrameric spectrin; in dimeric spectrin the self-association domain has not dimerized across two spectrin hetero-dimers to give the dimerized self-association domain. However available data indicates that the labile tail of the β-tetramerization domain forms a loop which mimics the same (56). In our present study neither the α-tetramerization domain nor the β-tetramerization domain could by themselves bind Prodan to a level that could be experimentally followed; only when they had dimerized to yield the reconstituted self-association domain could they bind Prodan. This can be explained by the fact that in both the spectrin dimer and tetramer the β-tetramerization domain’s labile tail is in a complex which forms a completed spectrin repeat domain, either by forming a loop or by complexing with a partial repeat domain in α-tetramerization domain (56-58) and it can be hypothesized that only when this complexed domain exists can Prodan bind. In the present work it is seen that the affinity of Prodan binding to the self-association domain is lower than that of binding to spectrin; this can be explained by the fact that while the α and β-tetramerization domains are capable of reconstituting the self-association domain, the structure of the same is not exactly the same as in the native condition, this is due to the fact that spectrin domains by themselves have less stability and structure than the whole protein (59).

The hydrophobic probe ANS was seen to bind to all the fragments of spectrin and spectrin itself. Generally native proteins do not bind ANS and ANS binding indicates some denaturation of a protein (60-62); however in cases where a protein has surface exposed hydrophobic patches the native protein can also bind ANS (63), making it a good probe for measuring protein surface hydrophobicity (64). We see that while all the domains bind ANS the affinity for the same is less than that in spectrin; moreover spectrin is shown to bind 5 ANS residues while the domains along with the self-association domain each bind 1 ANS for a total of 10. It is possible that intact spectrin has some hotspots of surface hydrophobicity which can bind ANS with much greater affinity thereby masking the binding of ANS to the other less-hydrophobic sites and the actual binding sites for ANS on spectrin is greater than 5. Present data shows us that all the fragments of spectrin have hydrophobic patches on their surface that can bind ANS and the hydrophobicity is of a comparable amount as evidenced by comparable dissociation constants for ANS binding.

The chaperone activity of spectrin fragments was assayed by following their ability to prevent protein aggregation and aid enzyme refolding. It was seen that in case of prevention of protein aggregation, intact spectrin was the best at protecting the aggregating proteins followed by the reconstituted self-association domain, all the other fragments had about the same activity which was slightly less than that of spectrin. It can be hypothesized that the spectrin fragments irrespective of their mutual structural differences act in a generalized manner when it comes to interacting with their chaperone substrates. It is plausible to think of the chaperone potential of spectrin fragments to be driven by the presence of surface exposed hydrophobic patches, as is the case with many chaperones (65,66), for example the chaperones such as sHSPs and GroEL also act with the help of exposed hydrophobic patches (67,68). It is known that chaperones act by recognizing the exposed hydrophobic clusters in structurally perturbed, misfolded, partially folded or stressed proteins with the help of hydrophobic patches of their own (29). As we have confirmed the presence of hydrophobic patches on all the spectrin domains and have determined that they all have about the same hydrophobicity it is reasonable to think that they are the determining factor in chaperone activity. Moreover the relatively unstable nature of the individual domains in comparison to intact spectrin (69,70) can be seen in the case of ADH aggregation where the difference in protection between spectrin and the cloned domains are most prominent probably due to the reduced stability of the domains in thermal stress. It is also interesting to note that spectrin and spectrin domains display a great specificity towards α-globin in favour of β-globin as a chaperone substrate as is evidenced by a large protection from aggregation of the former versus a modest one of the latter (21).

Our previous studies have shown that spectrin displays a selective affinity towards α-globin as a substrate as versus β-globin. Basic sequence alignment and crystal structure comparison reveals that the spectrin repeat domains share a similar fold and moderate homology to the α-globin chaperone AHSP (71). We have hypothesised that α-globin may act as a major client for the chaperone activity of spectrin based on both its selective binding and high affinity (21,72).

Enzyme refolding studies illuminate the fact that the mechanism of chaperone action is not the same for all client proteins; in case of alkaline phosphatase the addition of spectrin and spectrin domains give a larger reactivation yield versus self refolding while in case of α-glucosidase we see the opposite to be true. Chaperones are known to act by either helping a protein refold as in case of HSP70 (73) or they bind to an unfolded protein stopping aggregation until another chaperone can refold the protein as in case of α-crystallins (74). It would seem that depending on the nature of the client, spectrin is able to follow either one of these two routes.

From our present study it can be concluded that the molecular site of Prodan binding in native dimeric erythroid spectrin is located in the self-association domain and that the chaperone activity of spectrin is derived from the presence of many surface exposed hydrophobic patches which give its constituent domains comparable chaperone activity with the self association domain being moderately better in terms of preventing protein aggregation.

## Experimental procedures

pET151/D-TOPO plasmids containing the sequences of interest were acquired from Invitrogen; fluorescent probes Prodan (6-propionyl-2[dimethylamino]-naphthalene) and ANS (1-anilinonaphthalene-8-sulfonic acid) were also from Invitrogen; PMB, PMSF, EDTA, DTT, sodium hydrogen phosphate, NaOH, Tris, Sephadex G-100, Sepharose CL-4B, DEAE cellulose and CM cellulose were purchased from Sigma. Insulin from bovine pancreas, alcohol dehydrogenase from *S. cerevisiae*, alkaline phosphatase from *E. coli*, α-glucosidase from *S. cerevisiae*, 4-nitrophenyl-α-D-glucopyranoside and para-itophenylphosphate were purchased from Sigma. Prism Ultra pre-stained protein ladder (3.5 – 254 kDa) was purchased from Abcam. All water used for experiments was purified via a Millipore system.

### Isolation of human erythroid spectrin

Human blood samples were collected from healthy volunteers with proper informed consent. Blood samples were obtained from Ramkrishna Mission Seva Pratisthan Hospital, Kolkata, India, with informed written consent of the patients following the guidelines of the Institutional Ethical Committee as elaborated earlier (72,75,76)

Dimeric human erythroid spectrin was isolated from clean white RBC ghost membranes following protocol elaborated in earlier studies (77). Briefly, RBCs were pelleted by centrifugation, washed with PBS containing 5 mM sodium phosphate, 155 mM NaCl, 1mM EDTA, pH 8.0 and lysed in hypotonic lysis buffer containing 5mM sodium phosphate, 1mM EDTA and 20 µg/ml PMSF. Resultant RBC membranes were collected by centrifugation and contaminating hemoglobin was removed by washing with lysis buffer to yield clean white ghosts. Spectrin was removed from ghost membranes by 30 min incubation at 37°C in low salt spectrin removal buffer containing 0.2 mM sodium phosphate, 0.1 mM EDTA, 0.2 mM DTT, 20 µg/ml PMSF, pH 8.0. Spectrin was further purified by 30% ammonium sulphate precipitation from crude extract and resultant preparation was run through Sepharose CL-4B column to give final pure product. Purity of preparation was checked by 8% SDS PAGE analysis and concentration of preparation was determined from known O.D. of 10.7 at 280 nm for 1% spectrin solution (77). Spectrin preparation was stored at − 20°C for a maximum of one month.

### Isolation and purification of human Hemoglobin and constituent α and β chains

Human hemoglobin variant HbA was isolated from blood samples collected from healthy volunteers with proper informed consent. RBCs were collected by centrifugation, washed with PBS and lysed with three volumes of 1mM Tris-HCl pH 8.0 at 4°C and lysate was purified by gel filtration on Sephadex G-100 column (45×1cm) using 5mM Tris-HCl 50mM KCl pH 8.0 buffer. Purity was checked by 12% SDS PAGE analysis and concentration was checked using molar extinction coefficient of 125000 at 415 nm. Presence of oxy form of hemoglobin was confirmed using absorbance at 415 nm and 541 nm. Hemoglobin preparation was stored at − 70°C for a maximum of one week (78).

α and β globin subunits were isolated from hemoglobin using previously published protocol (78,79). Briefly, 100mg PMB per 1 g of hemoglobin was dissolved in minimum volume of 0.1M KOH and 1M acetic acid was added till very light precipitate persisted. PMB solution was added to 50mg/ml solution of hemoglobin in 10mM Tris-HCl 200mM NaCl, pH 8.0, and final pH was adjusted to 6.0 by addition of 1M acetic acid and mixture was incubated at 4° C for 12 hours.

PMB bound globin subunits were separated using two column selective ion exchange chromatography. α-PMB was isolated by equilibrating globin-PMB mixture with 10mM phosphate buffer pH 8.0 and passing through DEAE-cellulose column equilibrated with the same buffer. Similarly β-PMB was isolated by equilibrating globin-PMB mixture and passing through CM-cellulose column equilibrated with the same buffer. The PMB was removed from isolated globin chains by addition of 50 mM β-mercaptoethanol in 0.1M phosphate buffer pH 7.5. Globin chains were then purified by gel filtration on BioGel P2 column. The purity was checked by 12% SDS page analysis and concentration was measured by Bradford method using BSA as standard. Globin chains were stored at 4° C for not more than 48 hours (79).

### Expression isolation and purification of recombinant spectrin domains

Utilizing the work of Forget *et. al.* (4,14) the polypeptide sequence of the chosen domains was determined from the cDNA and polypeptide sequence of human erythroid spectrin (EMBL Data Bank accession numbers J05244 & J05500). In case of the EF domain which did not have any tryptophans in its sequence a tyrosine was replaced with a tryptophan in the 2578^th^ position in α-spectrin. pET151/D-TOPO plasmids with the sequences of interest inserted under control of Lac operon and T7 viral promoter with N-terminal hexa-histidine tag and Ampicillin selection marker was obtained from Invitrogen. Sequences were optimized for expression in *E. coli*. Plasmids were sequentially electroporated into XL1-Blue and BL21(DE3) cells for cloning and protein expression respectively.

It was noted from pilot experiments that the recombinant domains had appreciable solubility issues and were mostly incorporated into inclusion bodies. As such expression, extraction, purification and refolding methods were standardised for urea denatured extraction of the domains. Cells were grown in L.B. media with 100 µg/ml Ampicillin at 37 °C and protein expression was induced in a log phase culture, O.D._280nm_ 0.6, with 0.5 mM IPTG at 25 ° for four hours.

Cells were collected by centrifugation at 4000 x g and washed to remove excess media. Cells were lysed by sonication in a buffer containing 8 M urea, 10 mM Tris-HCl, 100 mM NaCl, pH 8.0 and lysate was clarified by centrifugation at 12000 x g to remove cell debris. Lysate was incubated with half its volume of Ni-NTA resin equilibrated with the same buffer for two hours at 25 °C. The resin was then washed successively with buffer and 20 mM immidazole in the same buffer to elute non-specific binders. Finally the proteins were eluted in 200 mM immidazole in the same buffer.

The resulting unfolded proteins were then sequentially dialyzed against 7 M, 6 M, 5 M, 4 M, 3 M, 2 M, 1 M and 0.5 M urea in the same buffer before being dialyzed in pure buffer without urea to yield folded protein. The resulting solution was centrifuged at 7000 x g for 5 minutes to remove any misfolded proteins and the supernatant was passed through 0.22 µm syringe filter to yield final pure folded proteins. The purity of the samples was checked by 15 % SDS-PAGE analysis and their concentration was measured using the Bradford method with BSA as standard. The proteins were stored at 4 °C for not more than 48 hours.

The self-association domain was reconstituted by incubating 20 µM concentrations of α and β-tetramerization domains together in 150 mM NaCl, 10 mM Tris-HCl pH 8.0 overnight on ice. Resulting reconstituted self-association domain was purified by passing it through a 45 x 1 cm Sephadex G-100 column equilibrated with 10 mM Tris-HCl, 100 mM NaCl, pH 8.0 buffer.

### Characterization of spectrin fragments

The fragments of spectrin were characterized using fluorescence and CD spectrometry both in native and urea denatured conditions. Steady state experiments were carried out in a Cary Varian and Fluoromax-3 spectrofluorometer using 10 mm path length quartz cuvettes at 25 °C using thermostated cuvette holders. 10 µM of each fragment and the reconstituted self-association domain were taken and their emission was measured from 310 to 450 nm using 295 nm excitation with 5nm band pass slits for both excitation and emission channels. The fluorescence anisotropy of the tryptophans in the proteins was measured using the same instrument with band pass slits of 5 nm each for excitation and emission and excitation at 295nm and emission at 340nm. Fluorescence lifetimes of these proteins were measured using a Horiba Fluoromax 3 with TCSPC attachment. Tryptophans were excited at 295 nm using a nano-LED and their emission was recorded at 340 nm. Decay curves were fitted using DAS6 software. CD spectroscopy was done using a 1 mm path length quartz sandwich type cuvette in a Applied Photophysics Chirascan instrument. An average of ten accumulations was taken for each sample using 0.5 nm steps in the interval of 190 to 250 nm. Machine data was converted to mean residual ellipticity for analysis and comparison.

### Binding of fluorescent probes

Prodan and ANS were used as fluorescent probes and binding experiments were performed using a Cary Varian spectrofluorometer using 10 mm path length quartz cuvettes at 25 °C using thermostated cuvette holders. Stock concentrations of Prodan in dimethylformamide were determined using an extinction coefficient of 18000 at 360 nm. Binding experiments were carried out in 10 mM Tris-HCl, 100 mM NaCl, pH 8.0 buffer, using 0.2 – 0.5 µM Prodan solutions to which increasing concentrations of proteins were added and emission was monitored in the range of 375 to 600 nm using an excitation of 360 nm and with 5 nm slits for both excitation and emission channels. The emission intensity at 430 nm (*I*_430_) and 520 nm (*I*_520_) was monitored for bound and free Prodan respectively (55).

Stock concentration of ANS in dimethylformamide was determined using an extinction coefficient of 7800 at 372 nm. Reverse titrations were done using 15 µM of protein in each case to which increasing concentrations of ANS were added. Fluorescence intensity was measured at 470 nm for bound ANS with excitation at 372 nm and band pass of 5 nm for both excitation and emission channels (20).

The extent of binding of ANS/Prodan was analyzed by a non-linear model independent method using the following equations to give the apparent dissociation constant in each case (K_d_).

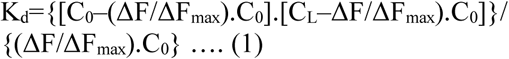

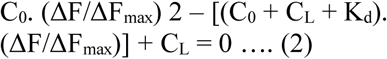

In equations (1) and (2), ΔF is the change in fluorescence emission intensity at 470 nm (for ANS) or the ratio of I_520_/I_430_ (for Prodan) for each point on the titration curve and ΔFmax denotes the same when a given probe is completely bound to a given protein, C_L_ is the concentration of the ligand protein being added at any given point in the titration curve, and C_0_ is the initial concentration of Prodan (opposite in case of ANS since reverse titration is done).

The y-axis intercept of the plot of 1/ΔF vs. 1/C_L_ gives the numerical value of 1/ ΔF_max_. In turn the calculated ΔF_max_ values can be put into equation 1 and the K_d_ values for each point on the titration curve can be generated, from which the mean K_d_ value is determined. The intersection of the tangents drawn on the initial and final linear region of the binding isotherm plot of ΔF/ΔF_max_ against C_L_ extrapolated to the ordinate gives the stoichiometry of Prodan binding to protein.

All experimental points for binding isotherm were fitted by least-square analysis using the Microcal Origin software package (Version 8.0) from Microcal Software Inc., Northampton, MA. The, K_d_ values are represented as mean ± standard error of the mean (S.D.) of at least 4 independent experiments (20).

The Scatchard equation was also used to estimate the intrinsic binding constants (K_o_) and the binding stoichiometry (n) according to the following equation.

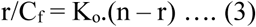

Where r = C_b_/C_p_, where C_b_ is the concentration of protein bound Prodan and C_p_ is the input concentration of protein. The concentration of bound Prodan was determined by normalizing the input concentration of Prodan with ΔF/ΔF_max_ and in the case of ANS it was determined as a function of the ΔF_max_ of 5 µM of ANS titrated against increasing concentrations of spectrin. The Scatchard plot was obtained by plotting r/C_f_ against r where C_f_ is defined as (C_o_ – C_b_) and C_o_ is the total concentration of Prodan. The best linear fit of the experimental data gave the values for K_o_ and n. Likewise ANS binding was similarly analysed.

In case of both Prodan and ANS the polarization and anisotropies were measured as a parameter for monitoring binding with steady state lifetimes being additionally monitored in case of Prodan.

### Assay of protein aggregation

Insulin, ADH, α and β globin were used as model substrates to check the chaperone potential of the spectrin domains.

A minimum volume of 20 mM NaOH was used to dissolve insulin which was then diluted to 0.3 mg/ml in 100 mM Na-phosphate buffer pH 7.0. Aggregation was induced by the addition of 25 µl of 1 M DTT to 1 ml of insulin solution and the aggregation was followed measuring the apparent increase in absorbance due to scattering at 360 nm as a function of time. Aggregation was carried out in the presence and absence of the spectrin domains and spectrin (20,22).

Similar aggregation experiments were carried out using 0.4 mg/ml of ADH in presence and absence of the spectrin domains and spectrin. Aggregation was induced by heating at 50 °C and followed by measuring the 90° light scattering at 450 nm.

Aggregation assays were carried out using α and β globin chains isolated from human hemoglobin using previously published protocol. Briefly, both kinds of globin chains were diluted with 50 mM Na-phosphate buffer pH 7.4, 150 mM NaCl, 1 mM EDTA to a final concentration of 13.5 μM and pre-incubated at 4 °C with spectrin or the spectrin domains for one hour. The samples were warmed to 37 °C and 10 μl of a 0.5 mM solution of potassium ferricyanide was added to induce aggregation. Aggregation was carried on in the presence and absence of spectrin or spectrin domains. Aggregation was monitored by measuring the apparent increase in absorbance due to scattering at 700 nm with time (21,80).

### Folding of denatured enzymes

Alkaline phosphatase in a final concentration of 500 µg/ml was denatured in 6 M guanidine-HCl in 10 mM Tris-HCl pH 8.0 buffer at 25 °C for an hour and then diluted 50 fold in 10 mM Tris-HCl pH 8.0 in presence and absence of spectrin and spectrin domains, to initiate refolding. Further 50 fold dilution was done in 0.5 mM Tris-HCl, pH 8.0 and para-nitophenylphosphate was added to a final concentration of 10 µM and incubated for 20 minutes. 100 mM final concentration of K_2_HPO_4_ was added to quench the reaction and absorbance was measured at 400 nm to monitor reactivation yield taking the activity of the native enzyme as 100% (20).

α-glucosidase in final concentration of 400 µg/ml was denatured in 8 M urea in 50mM Na-phosphate pH 7.6 for two hours. Refolding was initiated by 100 fold dilution in the same buffer in presence and absence of spectrin and spectrin domains. 0.3 mM final concentration of para-nitophenyl-α-D-glucopyranoside was used as enzyme substrate and the velocity of enzymatic reaction was followed by monitoring the increase in absorbance at 400 nm with time. Activity of native enzyme was taken as 100% (20).

## Abbreviations

DTT: di-thiothreitol
RBC: red blood cell
DEAE: diethylaminoethyl
CM: carboxymethyl
HbA: hemoglobin A
Prodan: 6-propionyl-2[dimethylamino]-naphthalene
ANS: 1-anilinonaphthalene-8-sulfonic acid
PMB: para-hydroxymercuribenzoic acid
ADH: alcohol dehydrogenase.

## References

1. Bennett, V., and Baines, A. J. (2001) Spectrin and ankyrin-based pathways: metazoan inventions for integrating cells into tissues. Physiological reviews 81, 1353–1392

2. Bennett, V., and Healy, J. (2009) Membrane domains based on ankyrin and spectrin associated with cell-cell interactions. Cold Spring Harbor perspectives in biology 1, a003012

3. Bennett, V. (1985) The membrane skeleton of human erythrocytes and its implications for more complex cells. Annual review of biochemistry 54, 273–304

4. Sahr, K. E., Laurila, P., Kotula, L., Scarpa, A. L., Coupal, E., Leto, T. L., Linnenbach, A. J., Winkelmann, J. C., Speicher, D. W., Marchesi, V. T., and et al. (1990) The complete cDNA and polypeptide sequences of human erythroid alpha-spectrin. The Journal of biological chemistry 265, 4434–4443

5. Pascual, J., Pfuhl, M., Walther, D., Saraste, M., and Nilges, M. (1997) Solution structure of the spectrin repeat: a left-handed antiparallel triple-helical coiled-coil. Journal of molecular biology 273, 740–751

6. Grum, V. L., Li, D., MacDonald, R. I., and Mondragon, A. (1999) Structures of two repeats of spectrin suggest models of flexibility. Cell 98, 523–535

7. Mirijanian, D. T., Chu, J. W., Ayton, G. S., and Voth, G. A. (2007) Atomistic and coarse-grained analysis of double spectrin repeat units: the molecular origins of flexibility. Journal of molecular biology 365, 523–534

8. An, X., Guo, X., Zhang, X., Baines, A. J., Debnath, G., Moyo, D., Salomao, M., Bhasin, N., Johnson, C., Discher, D., Gratzer, W. B., and Mohandas, N. (2006) Conformational stabilities of the structural repeats of erythroid spectrin and their functional implications. The Journal of biological chemistry 281, 10527–10532

9. Stabach, P. R., Simonovic, I., Ranieri, M. A., Aboodi, M. S., Steitz, T. A., Simonovic, M., and Morrow, J. S. (2009) The structure of the ankyrin-binding site of beta-spectrin reveals how tandem spectrin-repeats generate unique ligand-binding properties. Blood 113, 5377–5384

10. Andrade, M. A., Perez-Iratxeta, C., and Ponting, C. P. (2001) Protein repeats: structures, functions, and evolution. Journal of structural biology 134, 117–131

11. Speicher, D. W., and Marchesi, V. T. (1984) Erythrocyte spectrin is comprised of many homologous triple helical segments. Nature 311, 177–180

12. MacDonald, R. I., Musacchio, A., Holmgren, R. A., and Saraste, M. (1994) Invariant tryptophan at a shielded site promotes folding of the conformational unit of spectrin. Proceedings of the National Academy of Sciences of the United States of America 91, 1299–1303

13. Djinovic-Carugo, K., Gautel, M., Ylanne, J., and Young, P. (2002) The spectrin repeat: a structural platform for cytoskeletal protein assemblies. FEBS letters 513, 119–123

14. Winkelmann, J. C., Chang, J. G., Tse, W. T., Scarpa, A. L., Marchesi, V. T., and Forget, B. G. (1990) Full-length sequence of the cDNA for human erythroid beta-spectrin. The Journal of biological chemistry 265, 11827–11832

15. Bennett, V., and Gilligan, D. M. (1993) The spectrin-based membrane skeleton and micron-scale organization of the plasma membrane. Annual review of cell biology 9, 27–66

16. Bennett, V., and Lambert, S. (1991) The spectrin skeleton: from red cells to brain. The Journal of clinical investigation 87, 1483–1489

17. Machnicka, B., Grochowalska, R., Boguslawska, D. M., Sikorski, A. F., and Lecomte, M. C. (2012) Spectrin-based skeleton as an actor in cell signaling. Cellular and molecular life sciences : CMLS 69, 191–201

18. Deng, H., Wang, W., Yu, J., Zheng, Y., Qing, Y., and Pan, D. (2015) Spectrin regulates Hippo signaling by modulating cortical actomyosin activity. eLife 4, e06567

19. Chakrabarti, A., Bhattacharya, S., Ray, S., and Bhattacharyya, M. (2001) Binding of a denatured heme protein and ATP to erythroid spectrin. Biochemical and biophysical research communications 282, 1189–1193

20. Bhattacharyya, M., Ray, S., Bhattacharya, S., and Chakrabarti, A. (2004) Chaperone activity and prodan binding at the self-associating domain of erythroid spectrin. The Journal of biological chemistry 279, 55080–55088

21. Basu, A., and Chakrabarti, A. (2015) Hemoglobin interacting proteins and implications of spectrin hemoglobin interaction. Journal of proteomics 128, 469–475

22. Bose, D., Patra, M., and Chakrabarti, A. (2017) Effect of pH on stability, conformation, and chaperone activity of erythroid & non-erythroid spectrin. Biochimica et biophysica acta. Proteins and proteomics 1865, 694–702

23. Matsudaira, P. (1991) Modular organization of actin crosslinking proteins. Trends in biochemical sciences 16, 87–92

24. Pascual, J., Castresana, J., and Saraste, M. (1997) Evolution of the spectrin repeat. BioEssays : news and reviews in molecular, cellular and developmental biology 19, 811–817

25. Harper, S. Q., Crawford, R. W., DelloRusso, C., and Chamberlain, J. S. (2002) Spectrin-like repeats from dystrophin and alpha-actinin-2 are not functionally interchangeable. Human molecular genetics 11, 1807–1815

26. Kennedy, S. P., Warren, S. L., Forget, B. G., and Morrow, J. S. (1991) Ankyrin binds to the 15th repetitive unit of erythroid and nonerythroid beta-spectrin. The Journal of cell biology 115, 267–277

27. Bennett, V. (1989) The spectrin-actin junction of erythrocyte membrane skeletons. Biochimica et biophysica acta 988, 107–121

28. Trave, G., Pastore, A., Hyvonen, M., and Saraste, M. (1995) The C-terminal domain of alpha-spectrin is structurally related to calmodulin. European journal of biochemistry 227, 35–42

29. Bose, D., and Chakrabarti, A. (2017) Substrate specificity in the context of molecular chaperones. IUBMB life 69, 647–659

30. Haque, M. E., Ray, S., and Chakrabarti, A. (2000) Polarity estimate of the hydrophobic binding sites in erythroid spectrin: a study by pyrene fluorescence. Journal of fluorescence 10, 1–6

31. Ipsaro, J. J., Harper, S. L., Messick, T. E., Marmorstein, R., Mondragon, A., and Speicher, D. W. (2010) Crystal structure and functional interpretation of the erythrocyte spectrin tetramerization domain complex. Blood 115, 4843–4852

32. Shammas, S. L., Rogers, J. M., Hill, S. A., and Clarke, J. (2012) Slow, reversible, coupled folding and binding of the spectrin tetramerization domain. Biophysical journal 103, 2203–2214

33. Nicolas, G., Pedroni, S., Fournier, C., Gautero, H., Craescu, C., Dhermy, D., and Lecomte, M. C. (1998) Spectrin self-association site: characterization and study of beta-spectrin mutations associated with hereditary elliptocytosis. The Biochemical journal 332 (Pt 1), 81–89

34. Henniker, A., and Ralston, G. B. (1994) Reinvestigation of the thermodynamics of spectrin self-association. Biophysical chemistry 52, 251–258

35. Gallagher, P. G., Zhang, Z., Morrow, J. S., and Forget, B. G. (2004) Mutation of a highly conserved isoleucine disrupts hydrophobic interactions in the alpha beta spectrin self-association binding site. Laboratory investigation; a journal of technical methods and pathology 84, 229–234

36. Harper, S. L., Begg, G. E., and Speicher, D. W. (2001) Role of terminal nonhomologous domains in initiation of human red cell spectrin dimerization. Biochemistry 40, 9935–9943

37. Begg, G. E., Harper, S. L., Morris, M. B., and Speicher, D. W. (2000) Initiation of spectrin dimerization involves complementary electrostatic interactions between paired triple-helical bundles. The Journal of biological chemistry 275, 3279–3287

38. Li, D., Tang, H. Y., and Speicher, D. W. (2008) A structural model of the erythrocyte spectrin heterodimer initiation site determined using homology modeling and chemical cross-linking. The Journal of biological chemistry 283, 1553–1562

39. Li, D., Harper, S., and Speicher, D. W. (2007) Initiation and propagation of spectrin heterodimer assembly involves distinct energetic processes. Biochemistry 46, 10585–10594

40. Trave, G., Lacombe, P. J., Pfuhl, M., Saraste, M., and Pastore, A. (1995) Molecular mechanism of the calcium-induced conformational change in the spectrin EF-hands. The EMBO journal 14, 4922–4931

41. Musacchio, A., Noble, M., Pauptit, R., Wierenga, R., and Saraste, M. (1992) Crystal structure of a Src-homology 3 (SH3) domain. Nature 359, 851–855

42. Koch, C. A., Anderson, D., Moran, M. F., Ellis, C., and Pawson, T. (1991) SH2 and SH3 domains: elements that control interactions of cytoplasmic signaling proteins. Science 252, 668–674

43. Ipsaro, J. J., Huang, L., and Mondragon, A. (2009) Structures of the spectrin-ankyrin interaction binding domains. Blood 113, 5385–5393

44. Karinch, A. M., Zimmer, W. E., and Goodman, S. R. (1990) The identification and sequence of the actin-binding domain of human red blood cell beta-spectrin. The Journal of biological chemistry 265, 11833–11840

45. Branton, D., Cohen, C. M., and Tyler, J. (1981) Interaction of cytoskeletal proteins on the human erythrocyte membrane. Cell 24, 24–32

46. Viguera, A. R., Martinez, J. C., Filimonov, V. V., Mateo, P. L., and Serrano, L. (1994) Thermodynamic and kinetic analysis of the SH3 domain of spectrin shows a two-state folding transition. Biochemistry 33, 2142–2150

47. Pantazatos, D. P., and MacDonald, R. I. (1997) Site-directed mutagenesis of either the highly conserved Trp-22 or the moderately conserved Trp-95 to a large, hydrophobic residue reduces the thermodynamic stability of a spectrin repeating unit. The Journal of biological chemistry 272, 21052–21059

48. Chattopadhyay, A., Rawat, S. S., Kelkar, D. A., Ray, S., and Chakrabarti, A. (2003) Organization and dynamics of tryptophan residues in erythroid spectrin: novel structural features of denatured spectrin revealed by the wavelength-selective fluorescence approach. Protein science : a publication of the Protein Society 12, 2389–2403

49. Grantcharova, V. P., and Baker, D. (1997) Folding dynamics of the src SH3 domain. Biochemistry 36, 15685–15692

50. Ray, S., Bhattacharyya, M., and Chakrabarti, A. (2005) Conformational study of spectrin in presence of submolar concentrations of denaturants. Journal of fluorescence 15, 61–70

51. Kelkar, D. A., Chattopadhyay, A., Chakrabarti, A., and Bhattacharyya, M. (2005) Effect of ionic strength on the organization and dynamics of tryptophan residues in erythroid spectrin: a fluorescence approach. Biopolymers 77, 325–334

52. van Rooijen, B. D., van Leijenhorst-Groener, K. A., Claessens, M. M., and Subramaniam, V. (2009) Tryptophan fluorescence reveals structural features of alpha-synuclein oligomers. Journal of molecular biology 394, 826–833

53. Patra, M., Mukhopadhyay, C., and Chakrabarti, A. (2015) Probing conformational stability and dynamics of erythroid and nonerythroid spectrin: effects of urea and guanidine hydrochloride. PloS one 10, e0116991

54. Hill, S. A., Kwa, L. G., Shammas, S. L., Lee, J. C., and Clarke, J. (2014) Mechanism of assembly of the non-covalent spectrin tetramerization domain from intrinsically disordered partners. Journal of molecular biology 426, 21–35

55. Chakrabarti, A. (1996) Fluorescence of spectrin-bound prodan. Biochemical and biophysical research communications 226, 495–497

56. Harper, S. L., Li, D., Maksimova, Y., Gallagher, P. G., and Speicher, D. W. (2010) A fused alpha-beta “mini-spectrin” mimics the intact erythrocyte spectrin head-to-head tetramer. The Journal of biological chemistry 285, 11003–11012

57. Li, D., Harper, S. L., Tang, H. Y., Maksimova, Y., Gallagher, P. G., and Speicher, D. W. (2010) A comprehensive model of the spectrin divalent tetramer binding region deduced using homology modeling and chemical cross-linking of a mini-spectrin. The Journal of biological chemistry 285, 29535–29545

58. Speicher, D. W., DeSilva, T. M., Speicher, K. D., Ursitti, J. A., Hembach, P., and Weglarz, L. (1993) Location of the human red cell spectrin tetramer binding site and detection of a related “closed” hairpin loop dimer using proteolytic footprinting. The Journal of biological chemistry 268, 4227–4235

59. Menhart, N., Mitchell, T., Lusitani, D., Topouzian, N., and Fung, L. W. (1996) Peptides with more than one 106-amino acid sequence motif are needed to mimic the structural stability of spectrin. The Journal of biological chemistry 271, 30410–30416

60. Semisotnov, G. V., Rodionova, N. A., Razgulyaev, O. I., Uversky, V. N., Gripas, A. F., and Gilmanshin, R. I. (1991) Study of the “molten globule” intermediate state in protein folding by a hydrophobic fluorescent probe. Biopolymers 31, 119–128

61. Bolognesi, B., Kumita, J. R., Barros, T. P., Esbjorner, E. K., Luheshi, L. M., Crowther, D. C., Wilson, M. R., Dobson, C. M., Favrin, G., and Yerbury, J. J. (2010) ANS binding reveals common features of cytotoxic amyloid species. ACS chemical biology 5, 735–740

62. Christensen, H., and Pain, R. H. (1991) Molten globule intermediates and protein folding. European biophysics journal : EBJ 19, 221–229

63. Stryer, L. (1965) The interaction of a naphthalene dye with apomyoglobin and apohemoglobin. A fluorescent probe of non-polar binding sites. Journal of molecular biology 13, 482–495

64. Patra, M., Mitra, M., Chakrabarti, A., and Mukhopadhyay, C. (2014) Binding of polarity-sensitive hydrophobic ligands to erythroid and nonerythroid spectrin: fluorescence and molecular modeling studies. Journal of biomolecular structure & dynamics 32, 852–865

65. Reddy, G. B., Kumar, P. A., and Kumar, M. S. (2006) Chaperone-like activity and hydrophobicity of alpha-crystallin. IUBMB life 58, 632–641

66. Xu, W., Yuan, X., Xiang, Z., Mimnaugh, E., Marcu, M., and Neckers, L. (2005) Surface charge and hydrophobicity determine ErbB2 binding to the Hsp90 chaperone complex. Nature structural & molecular biology 12, 120–126

67. Haslbeck, M. (2002) sHsps and their role in the chaperone network. Cellular and molecular life sciences : CMLS 59, 1649–1657

68. Lin, Z., Schwartz, F. P., and Eisenstein, E. (1995) The hydrophobic nature of GroEL-substrate binding. The Journal of biological chemistry 270, 1011–1014

69. Lenne, P. F., Raae, A. J., Altmann, S. M., Saraste, M., and Horber, J. K. (2000) States and transitions during forced unfolding of a single spectrin repeat. FEBS letters 476, 124–128

70. MacDonald, R. I., and Pozharski, E. V. (2001) Free energies of urea and of thermal unfolding show that two tandem repeats of spectrin are thermodynamically more stable than a single repeat. Biochemistry 40, 3974–3984

71. Feng, L., Gell, D. A., Zhou, S., Gu, L., Kong, Y., Li, J., Hu, M., Yan, N., Lee, C., Rich, A. M., Armstrong, R. S., Lay, P. A., Gow, A. J., Weiss, M. J., Mackay, J. P., and Shi, Y. (2004) Molecular mechanism of AHSP-mediated stabilization of alpha-hemoglobin. Cell 119, 629–640

72. Mishra, K., Chakrabarti, A., and Das, P. K. (2017) Protein-Protein Interaction Probed by Label-free Second Harmonic Light Scattering: Hemoglobin Adsorption on Spectrin Surface as a Case Study. The journal of physical chemistry. B 121, 7797–7802

73. Bukau, B., and Horwich, A. L. (1998) The Hsp70 and Hsp60 chaperone machines. Cell 92, 351–366

74. Derham, B. K., and Harding, J. J. (1999) Alpha-crystallin as a molecular chaperone. Prog Retin Eye Res 18, 463–509

75. Chakrabarti, A., Bhattacharya, D., Deb, S., and Chakraborty, M. (2013) Differential thermal stability and oxidative vulnerability of the hemoglobin variants, HbA2 and HbE. PloS one 8, e81820

76. Basu, A., Saha, S., Karmakar, S., Chakravarty, S., Banerjee, D., Dash, B. P., and Chakrabarti, A. (2013) 2D DIGE based proteomics study of erythrocyte cytosol in sickle cell disease: altered proteostasis and oxidative stress. Proteomics 13, 3233–3242

77. Datta, P., Chakrabarty, S. B., Chakrabarty, A., and Chakrabarti, A. (2003) Interaction of erythroid spectrin with hemoglobin variants: implications in beta-thalassemia. Blood cells, molecules & diseases 30, 248–253

78. Datta, P., Chakrabarty, S., Chakrabarty, A., and Chakrabarti, A. (2007) Spectrin interactions with globin chains in the presence of phosphate metabolites and hydrogen peroxide: implications for thalassaemia. Journal of biosciences 32, 1147–1151

79. Datta, P., Basu, S., Chakravarty, S. B., Chakravarty, A., Banerjee, D., Chandra, S., and Chakrabarti, A. (2006) Enhanced oxidative cross-linking of hemoglobin E with spectrin and loss of erythrocyte membrane asymmetry in hemoglobin Ebeta-thalassemia. Blood cells, molecules & diseases 37, 77–81

80. Kihm, A. J., Kong, Y., Hong, W., Russell, J. E., Rouda, S., Adachi, K., Simon, M. C., Blobel, G. A., and Weiss, M. J. (2002) An abundant erythroid protein that stabilizes free alpha-haemoglobin. Nature 417, 758–763

