## supplementary material for "Localizing the chaperone activity of erythroid spectrin"

**Table ST1**

| Domain name | Domain location | Calculated weight | Sequence |
|---|---|---|---|
| α-Tetramerization | α-spectrin, 1-158 residues | 19 kDa | M E Q F P K E T V V E S S G P K V L E T A E E I Q E R R Q E V L T R Y Q S F K E R V A E R G Q K L E D S Y H L Q V F K R D A D D L G K W I M E K V N I L T D K S Y E D P T N I Q G K Y Q K H Q S L E A E V Q T K S R L M S E L E K T R E E R F T M G H S A H E E T K A H I E E L R H L W D L L L E L T L E K G D Q L L R A L |
| SH3 | α-spectrin, 973-1055 residues | 9.5 kDa | V E G V A G E Q R V M A L Y D F Q A R S P R E V T M K K G D V L T L L S S I N K D W W K V E A A D H Q G I V P A V Y V R R L A H D E F P M L P Q R R R E E P G N I T Q |
| Spectrin repeat | α-spectrin, 1470-1576 residues | 12 kDa | E I A T R L Q R V L D R W K A L K A Q L I D E R T K L G D Y A N L K Q F Y R D L E E L E E W I S E M L P T A C D E S Y K D A T N I Q R K Y L K H Q T F A H E V D G R S E Q V H G V I N L G N S L I E R S C D G N E E A |
| α-Dimerization | α-spectrin, 2002-2233 residues | 27 kDa | K A I E E R Y A A L L K R W E Q L L E A S A V H R Q K L L E K Q L P L Q K A E D L F V E F A H K A S A L N N W C E K M E E N L S E P V H C V S L N E I R Q L Q K D H E D F L A S L A R A Q A D F K C L L E L D Q Q I K A L G V P S S P Y T W L T V E V L E R T W K H L S D I I E E R E Q E L Q K E E A R Q V K N F E M C Q E F E Q N A S T F L Q W I L E T R A Y F L D G S L L K E T G T L E S Q L E A N K R K Q K E I Q A M K R Q L T K I V D L G D N L E D A L I L D I K Y S |
| EF | α-spectrin, 2257-2429 residues | 19 kDa | Q I Q A K D I K G V S E E T L K E F S T I **W** K H F D E N L T G R L T H K E F R S C L R G L N Y Y L P M V E E D E H E P K F E K F L D A V D P G R K G Y V S L E D Y T A F L I D K E S E N I K S S D E I E N A F Q A L A E G K S Y I T K E D M K Q A L T P E Q V S F C A T H M Q Q Y M D P R V E A I S L A M T T L A S P I P T L A T N K Q L L V D R R K S |
| Actin Binding | β- Spectrin, 1-136 residues | 36 kDa | M T S A T E F E N V G N Q P P Y S R I N A R W D A P D D E L D N D N S S A R L F E R S R I K A L A D E R E V V Q K K T F T K W V N S H L A R V S C R I T D L Y K D L R D G R M L I K L L E V L S G E M L P K P T K G K M R I H C L E N V D K A L Q F L K E Q R |

| | | | |
|---|---|---|---|
| | | | V H L E N M G S H D I V D G N H R L<br>V L G L I W T I I L R F Q I Q D I V V<br>Q T Q E G R E T R S A K D A L L L W<br>C Q M K T A G Y P H V N V T N F T S<br>S W K D G L A F N A L I H K H R P D<br>L I D F D K L K D S N A R H N L E H<br>A F N V A E R Q L G I I P L L D P E D<br>V F T E N P D E K S I I T Y V V A F Y<br>H Y F S K M K V L A V E G K R V G<br>K V I D H A I E T E K M I E K Y S G L<br>A S D L L T |
| β-Dimerization | β- Spectrin, 271-500 residues | 27 kDa | V A F Y H Y F S K M K V L A V E G K<br>R V G K V I D H A I E T E K M I E K<br>Y S G L A S D L L T W I E Q T I T V L<br>N S R K F A N S L T G V Q Q Q L Q A<br>F S T Y R T V E K P P K F Q E K G N<br>L E V L L F T I Q S R Met R A N N Q<br>K V Y T P H D G K L V S D I N R A W<br>E S L E E A G Y R R E L A L R N E L I<br>R Q E K L E Q L A R R F D R K A A M<br>R E T W L N E N Q R L V A Q D N F G<br>Y D L A A V E A A K K K H E A I E T<br>D T A A Y E E R V R A L E D L A Q E<br>L E K E N Y H D Q K R I T |
| Ankyrin Binding | β- Spectrin, 1657-1876 residues | 25 kDa | E G E Q I I R L Q G Q V D K H Y A G<br>L K D V A E E R K R K L E N M Y H L<br>F Q L K R E T D D L E Q W I S E K E<br>L V A S S P E M G Q D F D H V T L L<br>R D K F R D F A R E T G A I G Q E R<br>V D N V N A F I E R L I D A G H S E<br>A A T I A E W K D G L N E M W A D<br>L L E L I D T R M Q L L A A S Y D L<br>H R Y F Y T G A E I L G L I D E K H R<br>E L P E D V G L D A S T A E S F H R<br>V H T A F E R D V H L L G V Q V Q Q<br>F Q D V A T R L Q T A Y A G E K A E<br>A I Q N |
| β-Tetramerization | β- Spectrin, 1895-2137 residues | 28 kDa | R R T Q L V D T A D K F R F F S M A<br>R D L L S W M E S I I R Q I E T Q E R<br>P R D V S S V E L L M K Y H Q G I N<br>A E I E T R S K N F S A C L E L G E S<br>L L Q R Q H Q A S E E I R E K L Q Q<br>V M S R R K E M N E K W E A R W E<br>R L R M L L E V C Q F S R D A S V A<br>E A W L I A Q E P Y L A S G D F G H<br>T V D S V E K L I K R H E A F E K S T<br>A S W A E R F A A L E K P T T L E L<br>K E R Q I A E R P A E E T G P Q E EE<br>G E T A G E A P V S H H A A T E R T<br>S P V S L W S R L S SS W E S L Q P E<br>P S H P Y |

**Table ST1:** The polypeptide sequence, amino acid positions on the spectrin subunits and calculated molecular weight of the domains are given. In EF domain the tyrosine that was replaced by tryptophan is highlighted.

**Figure S1**

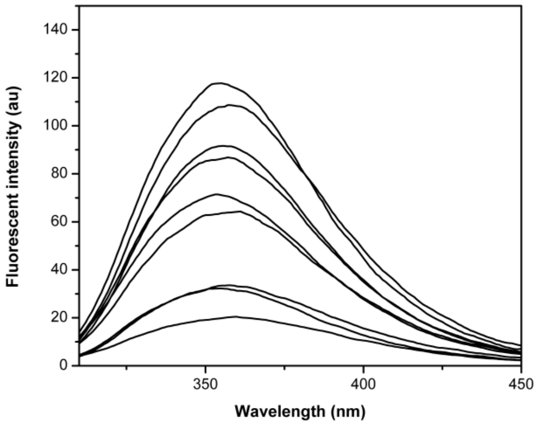


**Figure S1:** The emission spectra of the spectrin domains in 8 M urea denatured condition are shown. The emission maxima appear red shifted. In order from top to bottom the spectra are of the β-tetramerization, actin binding, α-dimerization, ankyrin binding, β-dimerization, spectrin repeat, SH3, α-tetramerization and EF domain respectively.

**Figure S2**

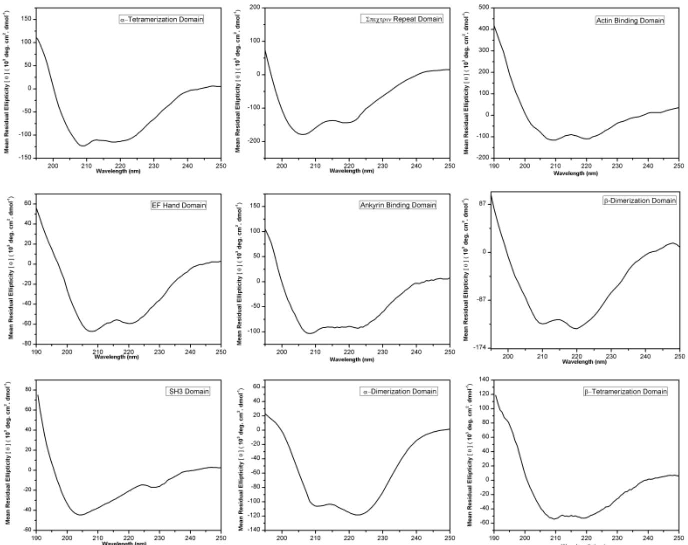


**Figure S2**: The CD spectra of the spectrin domains are shown in the range of 190 to 250 nm; spectra reveal a predominantly α-helical nature except SH3 domain which is globular.

**Figure S3**

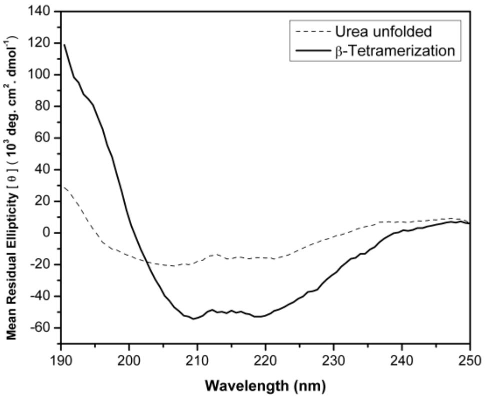


**Figure S3**: Representative CD spectra of β-tetramerization domain in 8 M urea denatured condition versus native condition are shown. Urea denatured condition leads to loss of secondary structure as evidenced by CD spectra from 190-250 nm.

**Figure S4**

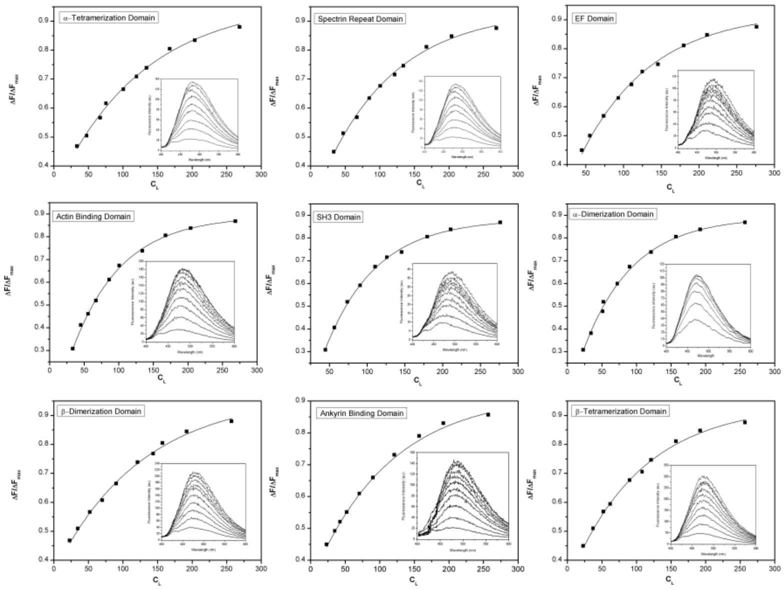


**Figure S4:** The binding isotherms generated by the model independent method for spectrin domain-ANS binding are shown; inset shows fluorescence spectra of the same.

**Figure S5**

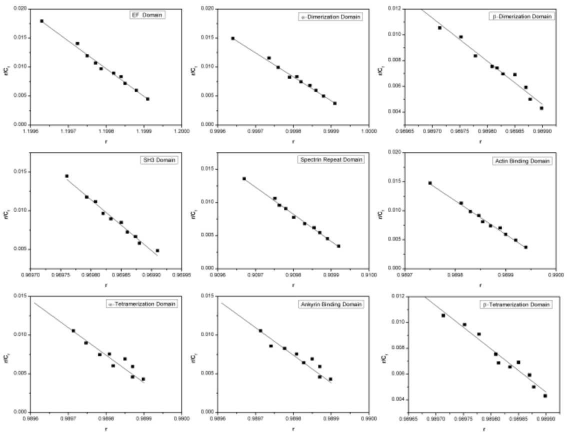


**Figure S5:** The Scatchard plots of ANS binding to spectrin domains are shown.
